# An atlas of inter- and intra-tumor heterogeneity of apoptosis competency in colorectal cancer tissue at single cell resolution

**DOI:** 10.1101/2021.03.19.436184

**Authors:** Andreas Ulrich Lindner, Manuela Salvucci, Elizabeth McDonough, Sanghee Cho, Xanthi Stachtea, Emer Patsy O’Connell, Alex D Corwin, Alberto Santamaria-Pang, Steven Carberry, Michael Fichtner, Sandra Van Schaeybroeck, Pierre Laurent-Puig, John P Burke, Deborah A McNamara, Mark Lawler, Anup Soop, John F Graf, Markus Rehm, Philip Dunne, Daniel B Longley, Fiona Ginty, Jochen HM Prehn

**Affiliations:** Department of Physiology and Medical Physics, Royal College of Surgeons in Ireland University of Medicine and Health Sciences, 123 St. Stephen’s Green, Dublin 2, Ireland; Centre of Systems Medicine, Royal College of Surgeons in Ireland University of Medicine and Health Sciences, 123 St. Stephen’s Green, Dublin 2, Ireland; GE Research, Niskayuna, NY 12309, USA; Centre for Cancer Research & Cell Biology, Queen’s University Belfast, 97 Lisburn Road, Belfast, BT9 7AE, Northern Ireland, UK; Department of Surgery, Royal College of Surgeons in Ireland University of Medicine and Health Sciences, 123 St. Stephen’s Green, Dublin 2, Ireland; Centre de Recherche des Cordeliers, INSERM, CNRS, Université de Paris, Sorbonne Université, USPC, Equipe labellisée Ligue Nationale Contre le Cancer, Paris, France; Beaumont Hospital, Beaumont Road, Dublin 9, Ireland; Institute of Cell Biology and Immunology, University of Stuttgart, Allmandring 31, 70569 Stuttgart, Germany

## Abstract

Cancer cells’ ability to inhibit apoptosis is key to malignant transformation and limits response to therapy. Here, we performed multiplexed immunofluorescence analysis on tissue microarrays with 373 cores from 168 patients, segmentation of 2.4 million individual cells and quantification of 20 cell lineage and apoptosis proteins. Ordinary differential equation-based modelling of apoptosis sensitivity at single cell resolution was conducted and an atlas of inter- and intra-tumor heterogeneity in apoptosis susceptibility generated. We identified an enrichment for BCL2 in immune, and BAK, SMAC and XIAP in cancer cells. ODE-based modelling at single cell resolution identified an enhanced sensitivity of cancer cells to mitochondrial permeabilization and executioner caspase activation compared to immune and stromal cells, with significant inter- and intra-tumor heterogeneity. However, we did not find increased spatial heterogeneity of apoptosis signaling in cancer cells, suggesting that such heterogeneity is an intrinsic, non-genomic property not increased by the process of malignant transformation.

## Introduction

Alterations in apoptosis signaling is key step in tumorigenesis(Hanahan and Weinberg, 2011). In many cases, cancer epithelial cells over time acquire alterations in their genome or epigenome that either result in an up-regulation of anti-apoptotic or a down regulation of pro-apoptotic proteins. Examples for such (epi)genomic alterations include promoter methylation and copy number alterations (Berdasco and Esteller, 2010; Mauro et al., 2015), while single point mutations in apoptosis-regulating genes are relatively rarely observed. Previous quantitative studies in solid tumor tissues found significant, but often complex differences in levels of individual anti-or pro-apoptotic proteins between different patients (Lindner et al., 2013; Lindner et al., 2017; Salvucci et al., 2017; Salvucci et al., 2019b). Predictions of individual patient’s apoptosis susceptibility is further complicated by the signaling redundancies in key apoptosis pathways, in particular the mitochondrial apoptosis pathway. Here, activation of either BAK or BAX is sufficient to induce mitochondrial outer membrane permeabilization (MOMP) (Kalkavan and Green, 2018), and this process is inhibited by a variety of different anti-apoptotic Bcl-2 family proteins including BCL2 itself, BCL(X)L and MCL1(Certo et al., 2006; Kalkavan and Green, 2018). Research into Bcl-2 family proteins and other apoptosis signaling proteins have resulted in the development and subsequent clinical approval of apoptosis sensitizers as anti-cancer agents. For example, Venetoclax is a selective BCL2 antagonist now used for the treatment of chronic lymphocytic leukemia, small lymphocytic lymphoma and acute myeloid leukemia which are characterized by strong BCL2 overexpression and dependency (Roberts et al., 2016). In context of solid tumors, the entry of apoptosis sensitizers into clinical practice has been relatively slow, a fact that is partially explained by the lack of gene mutations or pronounced over-or under-expression of individual apoptosis signaling proteins in solid tumor cells which could otherwise serve as stratification tools in clinical trials.

To overcome such limitations, various groups have developed computational models that describe apoptosis sensitivity on a systems level. BH3-only proteins are upstream initiators of the mitochondrial apoptosis pathway that are activated transcriptionally or post-translationally in response to stresses, such as DNA damage, genotoxic drugs, irradiation or withdrawal of trophic support. BH3-only proteins activate BAK and BAX directly, or activate these indirectly by binding to and neutralizing anti-apoptotic Bcl-2 proteins (Leber et al., 2007). BH3-peptide profiling has been successfully applied to predict outcome and responses to cancer therapeutics in solid cancers, however this technique requires fresh tissue (Ni Chonghaile et al., 2011). Other groups, including our own, have used gene expression or protein level (Reverse Protein Phase Array, RPPA) data of apoptosis-regulating genes from fresh-frozen or formalin-fixed tissues as input into deterministic signaling network models to estimate the intrinsic apoptosis sensitivity of individual tumors (Lindner et al., 2017; Salvucci et al., 2017).

Notwithstanding the successful application of these techniques in predicting chemotherapy responses and clinical outcome in cancer patients, the above techniques usually require a tissue homogenate to be analyzed. However, such “bulk” profiling results in the loss of not only important spatial information but also the precise cell-of-origin of the signals. It is feasible that some cancer cell populations in a given tumor are more resistant to therapy than other cancer cells, which is in line with evidence indicating the role of tumor heterogeneity in determining clinical outcome and responses to therapy (Fisher et al., 2013; Marusyk et al., 2012). Such resistant cell populations could give rise to more aggressive tumors on recurrence. Similarly, chemo- or radiation therapy not only affects tumor cells, but also cells in the tumor microenvironment such as immune cells; therefore, a higher apoptosis sensitivity of anti-tumor immune cells compared to cancer epithelial cells may be detrimental to patients.

To describe the extent of inter-individual and intra-tumor heterogeneity in apoptosis signaling, herein we employed an innovative multiplexed immunofluorescence imaging technique (Cell DIVE™), which is comprised of a repeated stain-image-dye-inactivation sequence using direct antibody-fluorophore conjugates, as well as a small number of primary antibodies from distinct species with secondary antibody detection (Gerdes et al., 2013), followed by single cell segmentation in a colorectal tumor tissue cohort. Using this method, we imaged 20 proteins and mapped quantities of the key members of the mitochondrial apoptosis pathway to 2.4 million individual cells (of which 1.6 million were colorectal tumor epithelial cells). This enabled us to calculate each individual cell’s apoptosis sensitivity through single cell systems modelling, and quantitatively describe inter- and intra-tumor heterogeneity of the mitochondrial apoptosis pathway within the different cell types that constitute a colorectal tumor.

To assess intrinsic apoptosis sensitivity of individual tumors, we had previously applied *‘averaged’* protein levels of tissue, but never single cell levels to our experimentally validated models, APOPTO-CELL (Huber et al., 2007; Rehm et al., 2006) and DR_MOMP (Lindner et al., 2013). Studying single cells’ with our apoptosis models is providing us with new insights into the mechanisms underlying apoptosis and treatment resistance.

## Results

### Multiplexed immunofluorescence imaging generates single cell profiles of mitochondrial apoptosis pathway proteins in 1.6 million individual colorectal tumor cells

To explore the levels of key proteins of the mitochondrial apoptosis pathways in colorectal cancer (CRC) tissue at the single cell level, we performed Cell DIVE™ multiplexing of nine pro-and anti-apoptotic proteins in regions of resected primary tumors in 355 tumor cores derived from 164 stage III CRC patients.

Apoptosis signaling protein selected for analysis included BCL2, BCL(X)L, MCL1, BAK and BAX which regulate the process of mitochondrial outer membrane permeabilization (MOMP), as well as PRO-CASPASE 9, PRO-CASPASE 3, XIAP and SMAC (DIABLO) which control the process of executioner caspase activation downstream of MOMP. For both processes, we previously devised and experimentally validated ordinary differential equation (ODE)-based, deterministic models, APOPTO-CELL (Huber et al., 2007; Rehm et al., 2006) and DR_MOMP (Lindner et al., 2013), that calculate the sensitivity of cancer cells to undergo mitochondrial apoptosis with high accuracy (Lindner et al., 2017; Salvucci et al., 2017), using quantities of the above 9 proteins as model input. Additional proteins selected for this study included cell identity markers (CD3, CD4, CD8, CD45, FOXP3, PCK26 and cytokeratin AE1), established markers of cell proliferation (KI67), antigen-presenting protein (HLA-A) and bioenergetics (GLUT1, CA9), as well as proteins used for cell segmentation analysis (Na+/K+-ATPase, cytokeratin AE1, PCK26, and S6).

We proceeded with multiplexed data acquisition of colon tumor tissue as follows (Figure 1A): 1-5) FFPE cores where formalin fixed paraffin embedded (FFPE) tissue microarrays first underwent antigen retrieval, followed by repeated cycles of protein staining, imaging and dye inactivation using cyanine dyes (Cy3 and Cy5) conjugated antibodies. DAPI staining and a background image was acquired in the beginning of each cycle for quality control and image processing. After rudimentary image processing (including illumination correction, distortion, stitching and registration) 6) we performed cell segmentation and single cell densitometry analysis. 7) We assessed the image quality of each core and removed 48 cores with insufficient quality. 8) We corrected possible batch effects between the five slides applying affine matrix transformations using an averaged distribution of protein intensities as reference for each protein marker. 9) Finally, we performed core and single cell analysis of the markers and performed model calculations within different cell populations.

**Figure 1.**
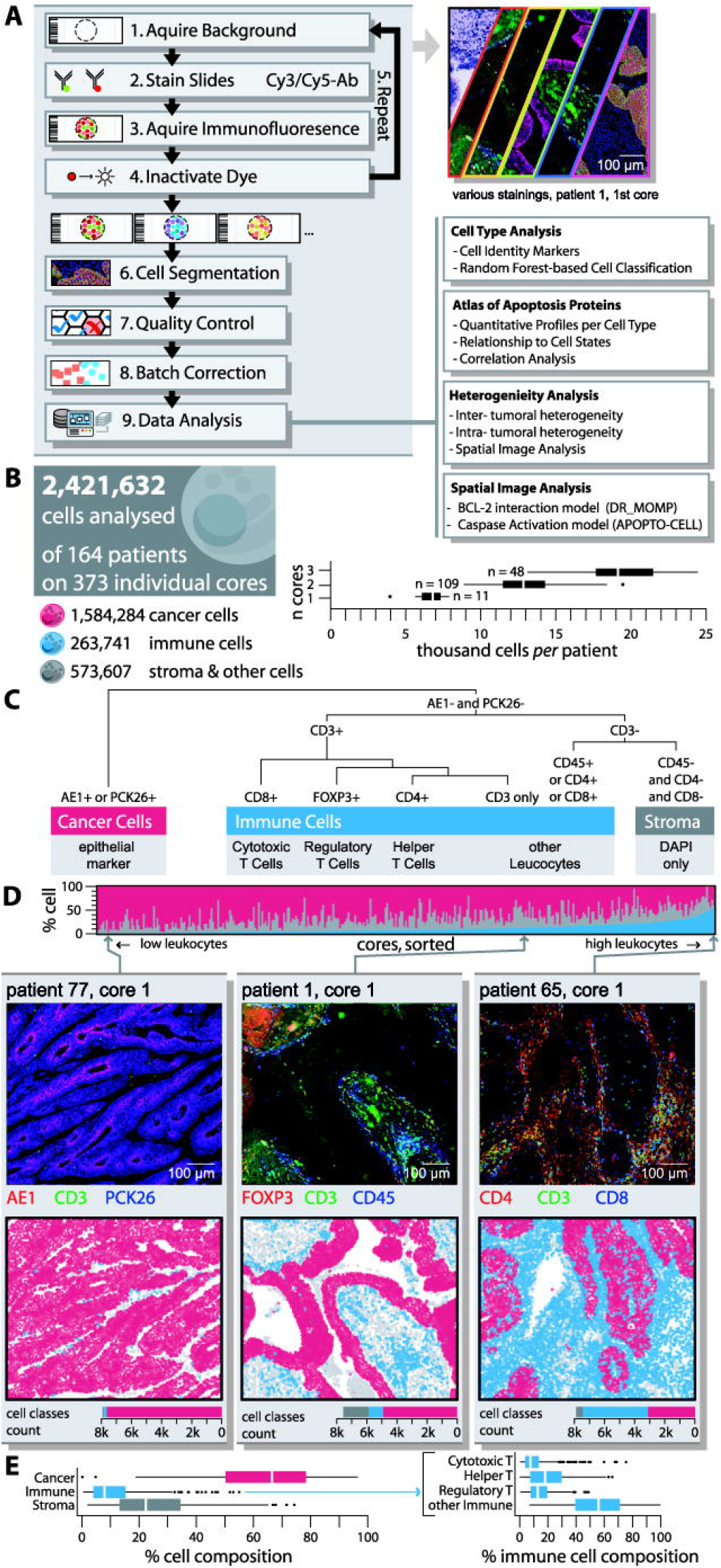
(A) Simplified workflow of the Cell DIVE™ platform and data analysis. (B) In total over 2 million cells, stratified into cancer, immune and stroma classes were analyzed. (C) Random forest was used to differentiate cells using DAPI, and epithelial and CD markers. (D) The majority of cores consisted of epithelial like cancer and stroma cells, (E) with less than 20% of cells being immune cells in the majority of cores (ANOVA, Tukey post-hoc).

This delivered a total of 54.6 million protein profiles (Figure 1B) in a total of 2.4 million cells which were used for cell identity analysis, construction of a tissue atlas of apoptotic proteins profiles, intra-and inter-tumor heterogeneity analysis, spatial tissue analysis as well as single cell systems modelling.

Cell DIVE™ and cell segmentation analysis identified on average 6,492 (SD 1,228) cells per tissue microarray (TMA) core; totaling on average 14,414 (SD 4,196) cells per patient (1 to 3 cores; Figure 1B). Cells were classified into different cell types based on cell identity markers for cancer/epithelial cells (positive for cytokeratin AE1 or PCK26), immune cells (positive for CD3, CD4, CD8 or CD45) and other stromal cells that were negative for any of these markers. For more extensive cell classification, a Random Forest model was trained with 15,184 manual annotated cells (0.6% of total cells) and CD3, CD4, CD8, CD45 and FOXP3, and applied on 99.9% of all cells to further differentiate immune cells into Cytotoxic, Regulatory, Helper T and other immune cells (Figure 1C). The model classified 65.7% as (epithelial like) cancer cells (type II error 3.0%; training set), 23.6% other stromal cells (type II error 8.1%) and 10.7% as immune cells (type II error 3.0%), of which 2.0% were Helper (type II error 28.8%), 1.4% Regulatory (type II error 7.4%), 1.3% Cytotoxic (type II error 28.0%) and 6.0% other T or immune cells (type II error 18.8%; Figure 1DE). Of note, the cell type composition in CRC core tissues varied significantly, with some cores showing predominantly cancerous/epithelial cells in the absence of immune cell infiltration, and others showing very high levels (up to 55%) of immune cells (Figure 1D). The median distribution of cells was 66.5% tumor, 7.8% immune and 22.2% stromal cells (Figure 1E). A bootstrap analysis with sampled pairings suggested that cell type composition in tumors of patients with paired-cores were, despite high heterogeneity, more similar to each other compared to random pairings. This suggests that cell type composition was a biological feature of individual tumors (Suppl. Figure 1).

### Tumor cell atlas shows heterogeneous and cell-type specific enrichment of key proteins of the mitochondrial apoptosis pathway

We next calculated molar protein profiles for proteins that are key to control MOMP and are used as input for the deterministic systems model, DR_MOMP. For the calculations of protein profiles of individual cells, we normalized cell intensities to the mean intensity in HeLa cells and used previously established concentrations in HeLa cells as reference (Flanagan et al., 2015; Lindner et al., 2013).

Analysis of the five key BCL2 proteins that control the process of MOMP demonstrated a significant enrichment in anti-apoptotic BCL2 in immune cells when compared to cancerous epithelial cells or other stromal cells, while anti-apoptotic BCL(X)L and MCL1 although statistically enriched in cancer epithelial cells were more homogenously distributed between the three cell types (Figure 2A-C). Mean levels of MCL1 were in general lower compared to BCL2 and BCL(X)L, confirming previous studies (Lindner et al., 2013). Of note, pro-apoptotic BAK showed a strong enrichment in cancer cells (Figure 2A-C), while again BAX, although statistically enriched in cancer cells, was more homogenously distributed between the three cell types.

**Figure 2.**
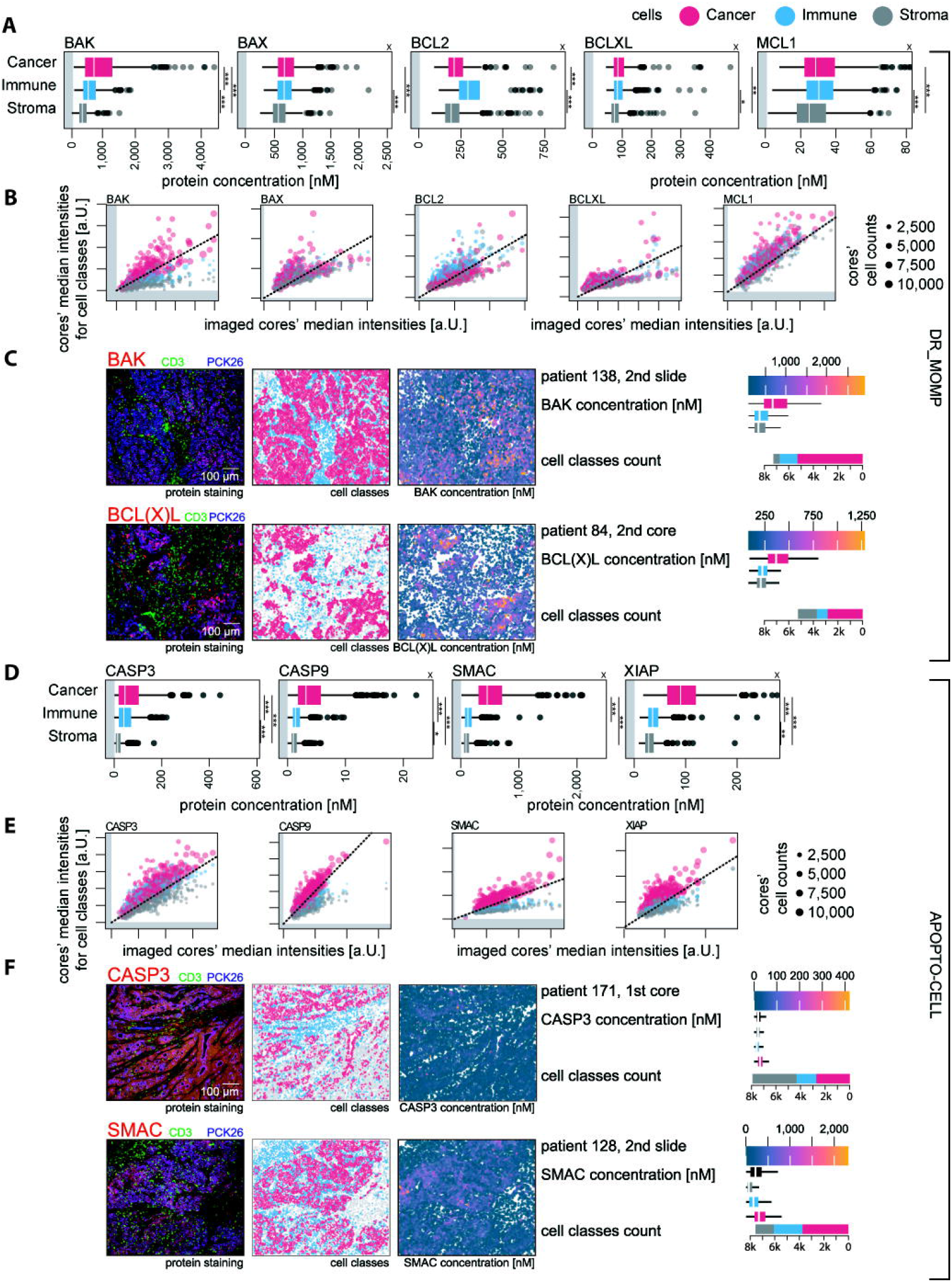
Protein analysis of apoptosis proteins relevant for (A-C) the DR_MOMP model upstream of MOMP and (D-F) the APOPTO-CELL model downstream of MOMP. (AD) To determine the difference between protein quantification based on cell masks and quantification using the whole image, we first determined the median protein concentration of each core, stratified for cancer (red), immune (blue) and stroma (gray) cells (ANOVA, Tukey post-hoc). *x* marks panels with cropped high value outliers. (BE) Subsequently, we compared the median pixel intensity of the core images (x-axes) with the stratified median pixel intensities determined using cell masks (y-axes) before batch correction. The scatter size indicates the numbers of stratified cells of the respective core. The panels C and F show examples of the pre-batch corrected protein staining, cell type classification and batch corrected mean cell intensities using cell masks.

For PRO-CASPASE 3, PRO-CASPASE 9, SMAC and XIAP single protein profiling we converted the batch-corrected protein intensities to µM concentrations via alignment with reference distributions (Hector et al., 2012) using a pipeline that we previously developed (Salvucci et al., 2019a; Salvucci et al., 2017). Proteins that control executioner caspase activation downstream of MOMP also showed a heterogeneous distribution between cell types, with XIAP, SMAC, PRO-CASPASE 3 and PRO-CASPASE 9, all at higher levels in cancer cells when compared to immune cells (Figure 2D-F). Stromal cells showed the lowest levels of these proteins, suggesting that the apoptotic machinery downstream of MOMP is suppressed in non-transformed cells when compared to cancer epithelial cells.

Utilizing transcriptional data derived from flow-sorted immune (n = 6), epithelial (n = 6) and fibroblast (n = 6) populations isolated from CRC primary tumor tissue (GSE39396 (Calon et al., 2012); Suppl. Table 2), we identified elevated levels of *bcl2* mRNA levels in leukocytes compared to cancer (epithelial) cells (ANOVA p = 0.006, Tukey post-hoc p = 0.005) but also significantly higher levels of bax and mcl1 mRNA levels in Leukocytes compared to cancer cells, and in Stroma (Fibroblasts) compared to cancer cells (ANOVA p ≤ 0.01, Tukey post-hoc p < 0.01; Suppl. Figure 2). We did not find any significant differences in mRNA levels of the *bak1, bcl2l1* (BCL(X)L), caspases, nor *xiap* between the cell populations.

Apoptotic protein profiles from approximate 115,923 identified T cells showed higher levels of BAK, XIAP, SMAC, PRO-CASPASE 3 and PRO-CASPASE 9 and lower levels of BCL2 in Cytotoxic T cells when compared to Helper or Regulatory T cells (Figure 3A-C). These findings suggests that Cytotoxic T cells may represent the T cells most sensitive to the activation of mitochondrial apoptosis.

**Figure 3.**
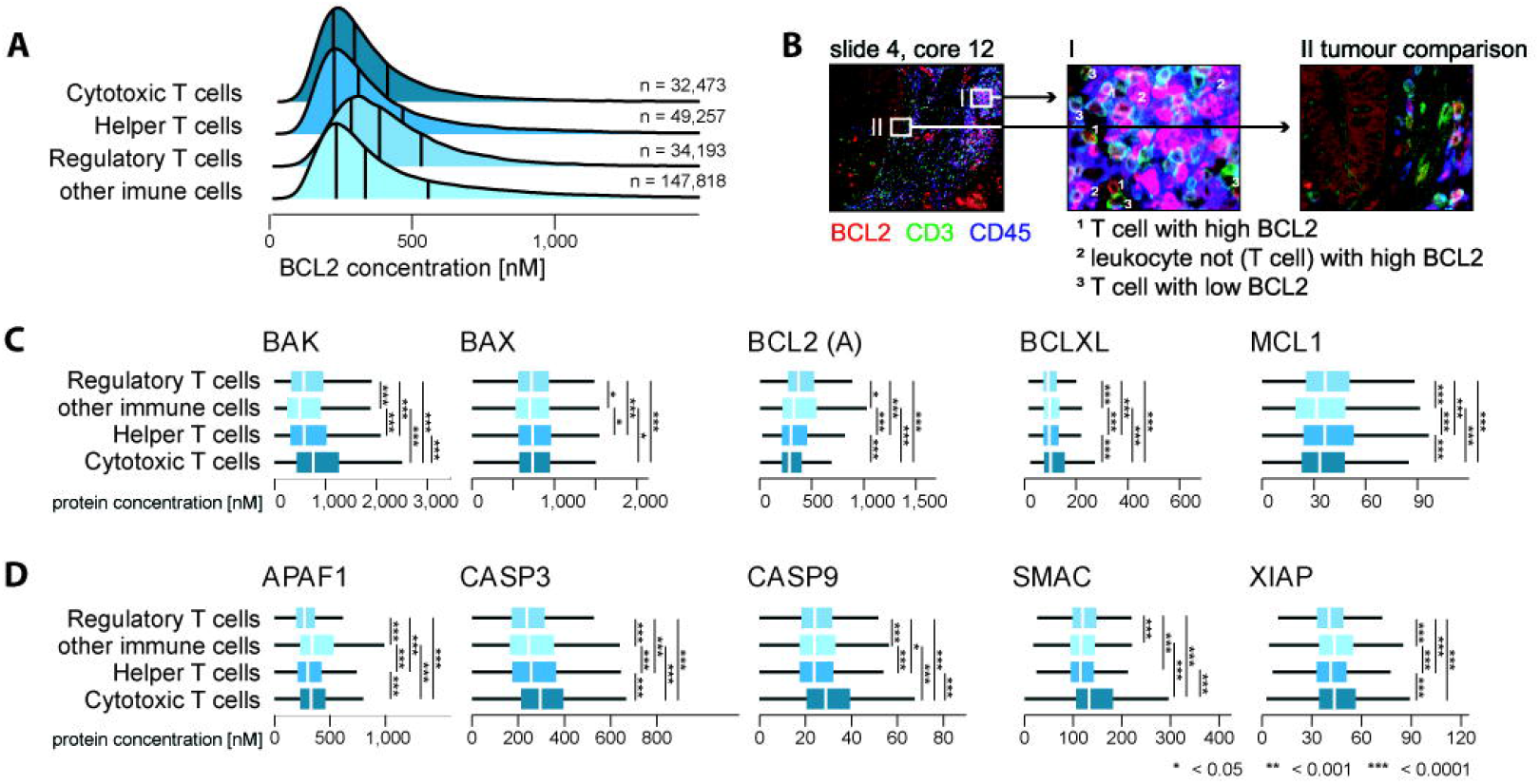
Global immune cell protein analysis of apoptosis proteins relevant for (A-C) the DR_MOMP model upstream of MOMP and (D) the APOPTO-CELL model downstream of MOMP (ANOVA, Tukey post-hoc). (B) Virtual IHC staining with BCL2 (red), CD3 (green) and CD45 (blue) shows that BCL2 level vary largly between immune cells.

As expected, cancer epithelial cells also showed higher levels of the glucose transporter GLUT1, sodium-potassium ATPase, the hypoxia-inducible factor-1α (HIF-1a) target gene CA9, and the proliferation marker KI67, while HLA-A were enriched in immune cells (Figure 4AB). In contrast, protein levels of p70S6 kinase (S6) were more evenly distributed across all cell types. Calculating the cores’ quartile coefficients of dispersion (COD; Suppl. Figure 3), a measure of the spread of the protein levels, we found that immune cells had a greater COD for BCL2 and BAK compared to cancer and stroma cells. Stroma cells showed the highest, and cancer epithelial cells the lowest, COD for MCL1, APAF1 and PRO-CASPASE 3. Cancer cells showed greater CODs of SMAC, GLUT1 and KI67 protein levels compared to immune and stroma cells.

**Figure 4.**
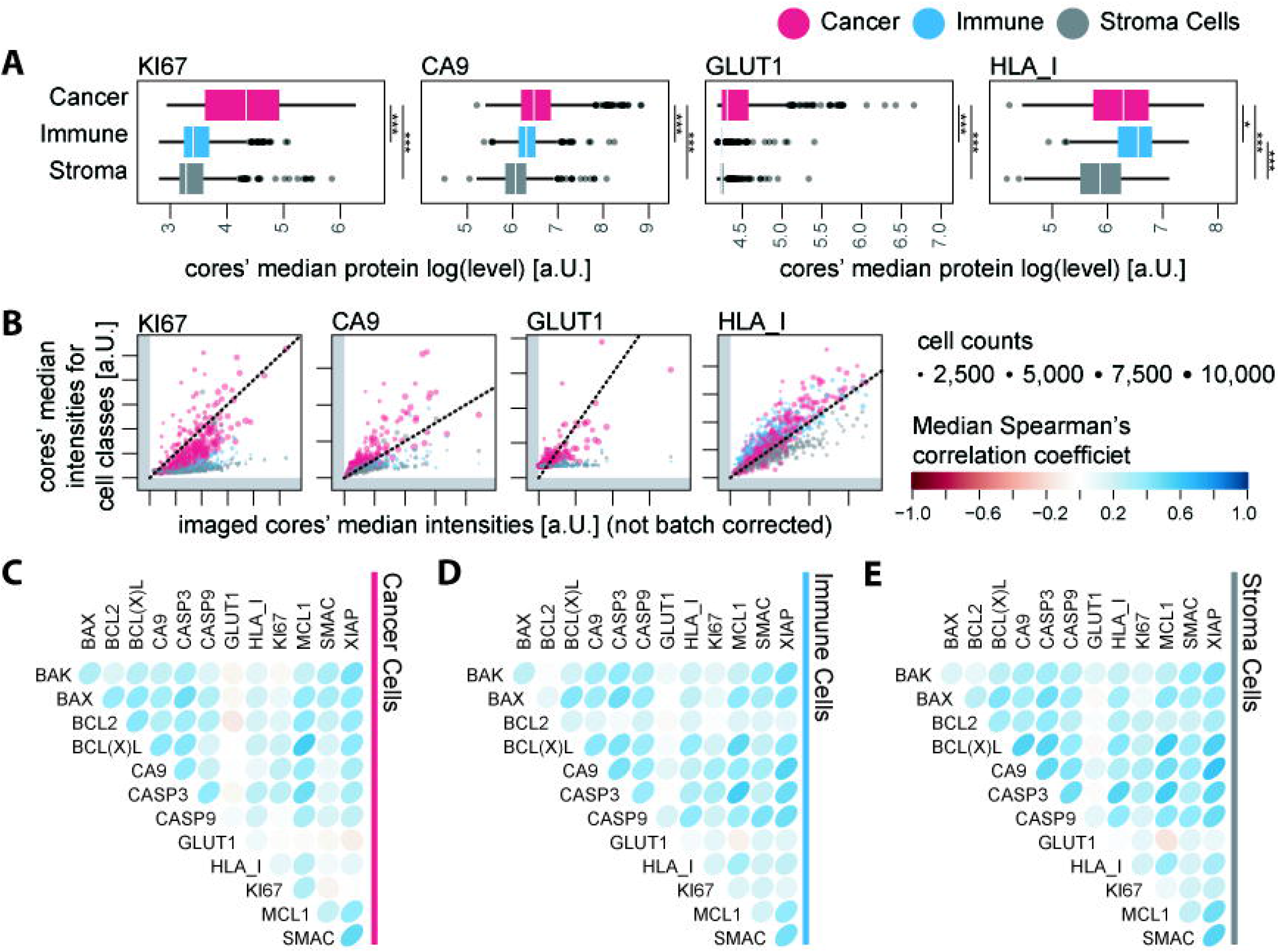
(A) Protein analysis of KI67, CA9, GLUT1 and HLA_I proteins using core median protein levels and stratified for cancer (red), immune (blue) and stroma (gray) cells (ANOVA, Tukey post-hoc). (B) We compared the median pixel intensity of the core images (x-axes) with the stratified median pixel intensities determined using cell masks (y-axes) before batch correction. The scatter size indicates the numbers of stratified cells of the respective core. We calculated the median spearman correlation coefficient between proteins, stratified for (C) cancer, (D) immune and (E) stroma cells. A more detailed correplation plot, including inter quantile ranges, is provided as supplementary figure 4.

Correlation analysis (Figure 4C-D) of the 1,556,581 cancer cells demonstrated high, positive median Spearman’s correlation coefficients (ρ > 0.5) between BAK and BAX levels. Levels between BAK (and BAX) and PRO-CASPASE 3 (and PRO-CASPASE 9), BCL(X)L and BCL2, PRO-CASPASE 3 and BCL2, BCL2 and MCL1, BCL2 and XIAP, SMAC and BCL(X)L, PRO-CASPASE 3 and PRO-CAPSASE 9, and PRO-CASPASE 3 and XIAP had high positive median correlation coefficients in cancer and stromal, but not immune cells. The Spearman’s correlation coefficient between BCL(X)L and MCL1, CA9 and XIAP, and SMAC and XIAP levels was > 0.5 in all cells. Comparing GLUT1 to apoptosis protein levels returned coefficients around 0, but showed greater values when compared to HLA-A and CA9 in cancer cells. HLA-A levels correlated with PRO-CASPASE 3 levels only in stromal cells. Generally, correlations between individual proteins were nearly identical in leukocytes and stromal cells and frequently differed from those in cancer cells, validating at the single cell level that transformed cells deviate from a physiological regulation of apoptotic and metabolic pathways.

### Single cell systems modelling of apoptosis sensitivity shows inter-individual differences in apoptosis sensitivity and an enhanced ability of tumor cells to undergo Caspase-3-dependent mitochondrial apoptosis

Next, we used quantitative single cell protein profiles to predict the apoptosis sensitivity of the 1.6 million colorectal tumor cells. We employed two systems models of the mitochondrial apoptosis pathway that were previously established and experimentally validated in our group to predict the intrinsic ability of cells to initiate and execute apoptosis. DR_MOMP (Flanagan et al., 2015; Lindner et al., 2013; Lindner et al., 2017) calculates the sensitivity of cells to undergo mitochondrial permeabilization by computing a score that, in summary, quantifies the amount of pro-apoptotic BH3-only proteins required to trigger sufficient BAK or BAX pore formation to induce mitochondrial outer membrane permeabilization (MOMP) during genotoxic stress (Figure 5A). In contrast, APOPTO-CELL (Huber et al., 2007; Rehm et al., 2006) calculates the amount of caspase 3 mediated substrate cleavage as a consequence of MOMP and apoptosome formation (Figure 5A).

**Figure 5.**
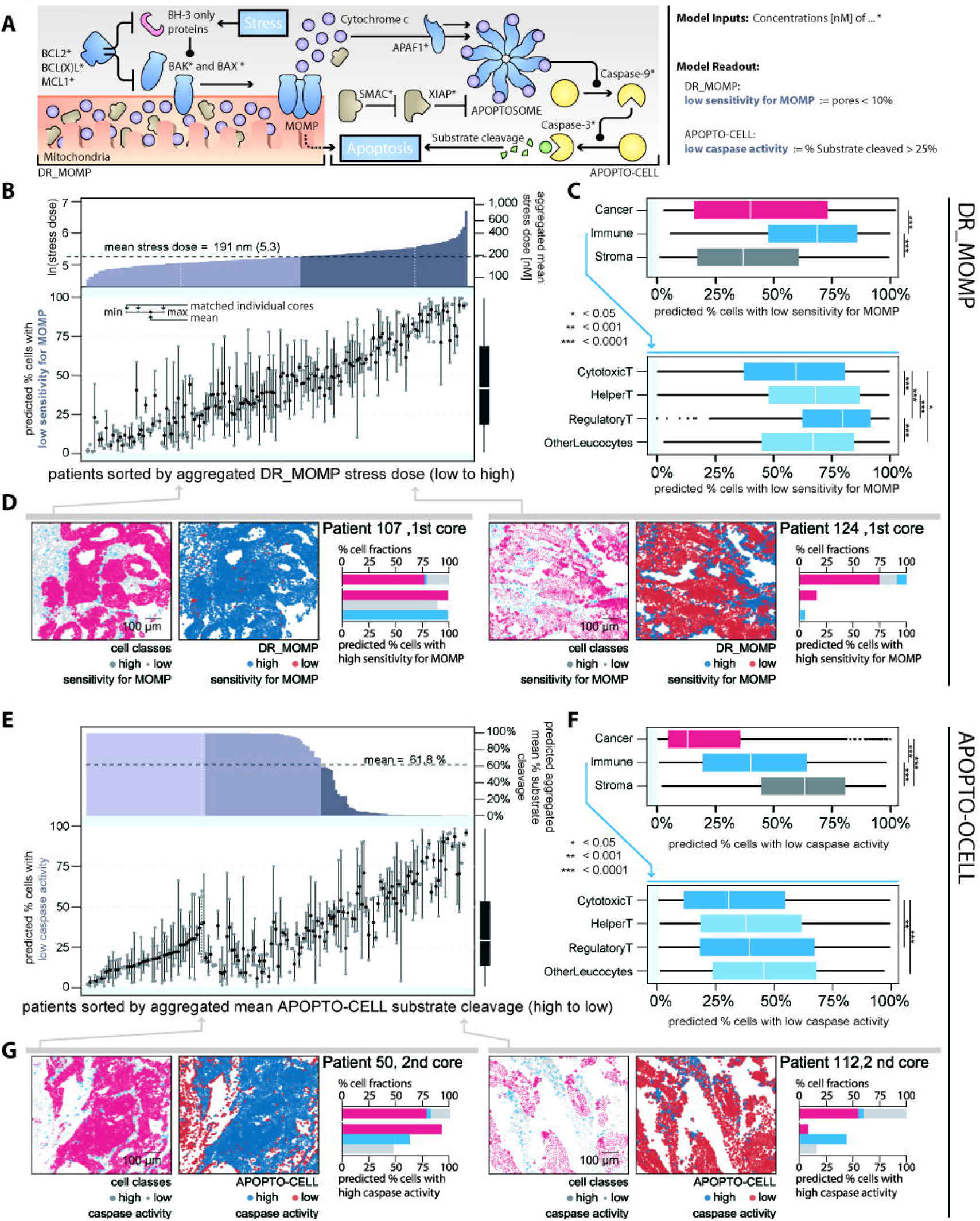
Results of the cell-by-cell analysis using the apoptosis models DR_MOMP (Lindner et al., 2013) and APOPTO-CELL (Huber et al., 2007; Rehm et al., 2006). (A) Graphical illustration of the modelled BCL2 pathway (DR_MOMP) upstream of MOMP and the modelled caspase pathway (APOPTO-CELL) downstream of MOMP. We first analyzed (B-D) DR_MOMP and subsequently (D-G) APOPTO-CELL. (BE) First we determined model predictions of required stress to induce MOMP (DR_MOMP) and % substrate cleavage upon MOMP (APOPTO-ELL) based on aggregated mean protein level for each patient, using the pool of all cells of multiple cores. Subsequently we calculated the cores’ cell fractions with (B) high/low sensitivity for MOMP (DR_MOMP) and (E) high/low substrate cleavage (APOPTO-CELL) using individual cell protein levels. We compared cores’ fractions with high/low (C) sensitivity for MOMP and (F) caspase activity stratified for cancer (red), immune (blue) and stroma (gray) cells (ANOVA, Tukey post-hoc).The panels D and G show examples of individual cores with high/low (D) sensitivity for MOMP and (G) caspase activity. In B and E, cores were sorted from high apoptosis sensitivity (left) to low apoptosis sensitivity (right), respectively.

Using the quantitative Bcl-2 protein profiles of BAK, BAX, BCL2, BCL(X)L and MCL1 as model input for DR_MOMP, we were able to calculate the sensitivity of individual tumor cells to the process of mitochondrial apoptosis initiation. Calculating mean DR_MOMP scores for each core (Figure 5B, top) and % cells with low sensitivity for MOMP for individual core (Figure 5B, bottom) using the calculated average stress dose of the population as threshold (Flanagan et al., 2015; Lindner et al., 2013; Lindner et al., 2017), we were able to show significant differences in % cells with low sensitivity for MOMP in this otherwise homogeneous cohort of stage III CRC patients. Between patient-matched cores, we found a mean difference of 18.8% ± SD 14.1% and a mean SD of 14.0% cells with low sensitivity for MOMP. When stratifying DR_MOMP calculations for individual cell types, we found that, on average, significantly fewer cancer cells and stromal cells exhibited low sensitivity for MOMP when compared to immune cells (Figure 5C, upper). Among immune cells, Regulatory T cells were found to have largest population of single cells with low sensitivity for MOMP (Figure 5C lower). And, in line with our analysis on protein level (Figure 3), we found that cytotoxic T cells are overall significantly more susceptible to apoptosis stimuli compared to other immune cells. Figure 5D depicts examples of DR_MOMP predictions in cores with a majority of cells having a high sensitivity for MOMP (left) or a majority of cells having low sensitivity for MOMP (right).

When investigating the sensitivity of individual tumor cells to undergo caspase 3 activation (once the process of MOMP is activated) using the APOPTO-CELL systems model, we similarly found significant differences between individual patients (Figure 5E, top) and cores (Figure 5E, bottom): Between patient-matched cores we found a mean difference of 18.8% ± SD 15.7% and a mean SD of 13.8% cells with low predicted caspase activity. Importantly, when investigating individual cell types, we found that cancer cells were predicted to show a higher caspase activity compared to immune cells and stromal cells, with the latter showing the greatest fraction of cells with low predicted caspase activity (Figure 5F). Figure 5G depicts examples of cores with APOPTO-CELL predicting the majority of cells showing high caspase activity (left) or the majority of cells exhibiting low caspase activity (right).

The activation of mitochondrial (or intrinsic) apoptosis is considered to be a two-step process, with little feed-back from one to the other process (Ichim and Tait, 2016). The multiplexing, quantitative protein profiling and single cell systems modelling pipeline developed here hence also allowed us to address the question of whether cancer cells show differences in their ability to activate each of these two apoptotic control points.

In line with the latter analysis, assessing apoptosis sensitivity up-and downstream of MOMP showed that cancer cells are sensitive for both apoptosis pathways in the majority of tumors and that only a small fraction of cores showed low sensitivity in both pathways at the same time (Figure 6A). In contrast, immune and stroma cells had a higher fraction of cells that showed low sensitivity in both pathways, and a lower fraction of cells that showed high sensitivity in both, compared to cancer cells (Figure 6AB). Of the cancer of cells that show low sensitivity in one and high sensitivity in the other pathway, we found that a majority of cancer cells showed a low MOMP sensitivity and a predicted high caspase activity (Figure 6A-C). In contrast, the majority of immune cells showed a predicted high caspase activity but a low sensitivity for MOMP (Figure 6A-C), and the majority of stroma cells showed a high sensitivity for MOMP but a predicted low caspase activity (Figure 6A-C). Collectively, the data suggested that the majority of cancer cells showed a high sensitivity for at least one of the two apoptosis pathways, and that cancer cells were overall more likely to respond to both signaling pathways when compared to immune or stromal cells.

**Figure 6.**
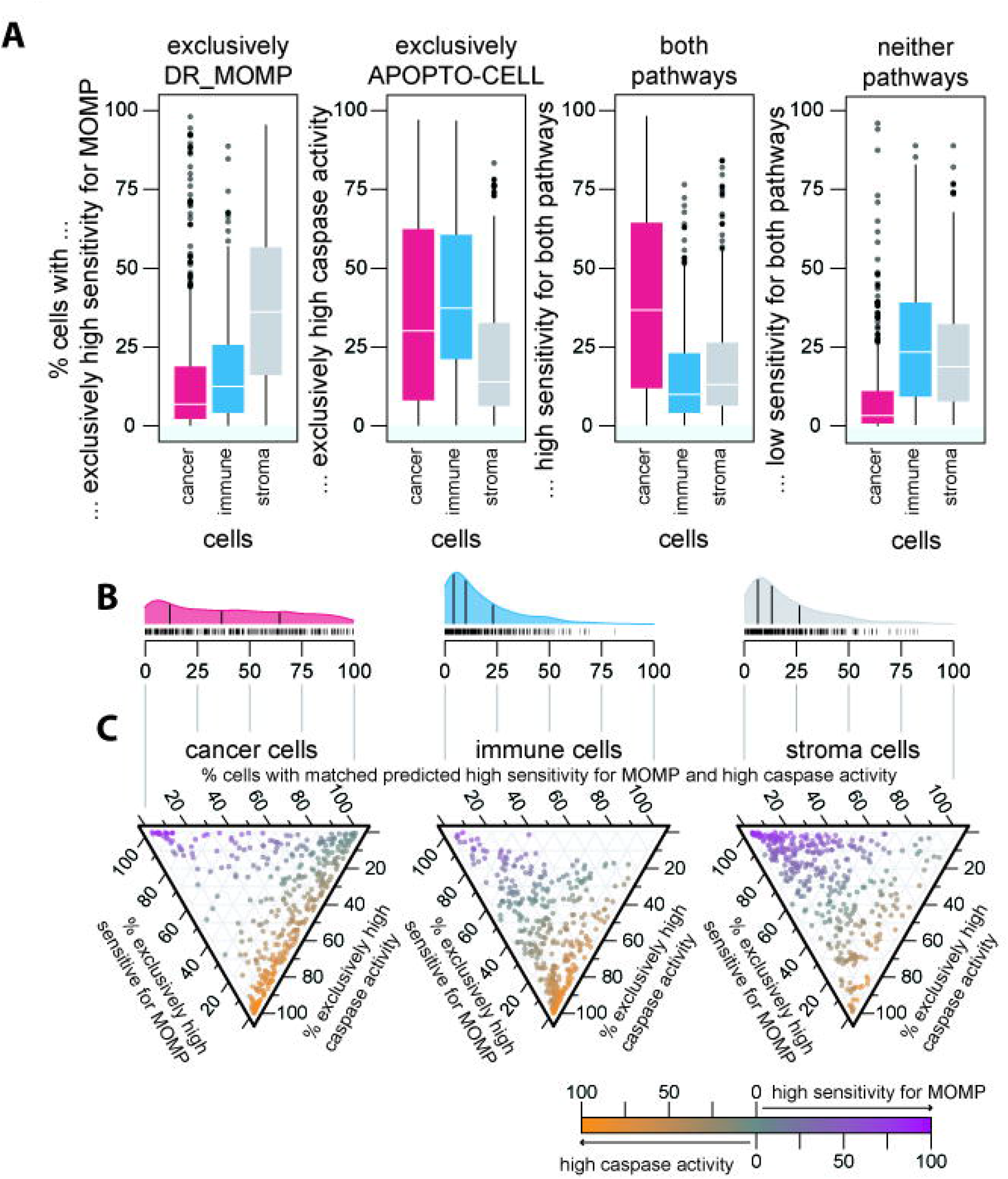
We determined cores’ cells that (A) exclusively showed high sensitivity for MOMP (left), high caspase activity, high responses in both apoptosis pathways and low responses in both apoptosis pathways (right; ANOVA, Tukey post-hoc).(BC) Ternary plot of individual core’s cell fraction for exclusively pathway responses or sensitivity in both pathways. Overall, cancer cells show high sensitivity for the DR_MOMP modelled BCL2 pathway upstream of MOMP with about half showing also high caspase activity modelled by APOPTO-CELL. Stroma cells showed exclusively high sensitivity for the apoptosis pathway upstream for MOMP while immune cells showed exclusively high sensitivity for MOMP.

### Analysis of intra-tumoral heterogeneity

While investigating apoptosis sensitivity at the single cell level using our systems models, we also noticed that certain patients showed a significant intra-tumor heterogeneity among cancer cells, while other patients showed a more homogenous distribution in model predictions (Figure 5BE).To further investigate such intra-tumoral heterogeneity, we assessed the Shannon Entropy between the models in each core to measure the unanimity of single-cell predictions. A low entropy, close to zero, suggests homogenous model predictions among all cells, which could either indicate systemic sensitivity or systemic resistance. In contrast, higher values suggest a more heterogeneous, or random, configuration of cell states, indicating a high diversity in distinct cells populations (Figure 7A). Overall, we did not find a significant difference in model predictions with the majority of cores having high entropy (> 0.5) for both models (Figure 7A). However, the cell composition of different cores may bias the calculation if not stratified for cell types. While the difference was small, cancer cells and immune cells had significantly lower entropy compared to stroma cells for DR_MOMP (Figure 7B). We found something similar for the predictions of the APOPTO-CELL model, however, the difference between cancer and immune cells was much more distinct (Figure 7C). Studying the entropy of protein levels using histograms (normalized bin size = 0.1 SD) in cancer cells (Figure 7D), suggested the highest entropy in BCL2 and the lowest entropy in MCL1 if comparing protein relevant for DR_MOMP. Among proteins relevant for APOPTO-CELL, we found the highest entropy in XIAP and the lowest PRO-CASPASE 3. Overall, cancer epithelial cells showed higher entropy in levels of all proteins but BCL(X)L and PRO-CASPASE 3 when compered between epithelial cancer, immune and stroma cells (ANOVA p < 0.05, Tukey post-hoc p < 0.05; Suppl. Figure 5). Figure 6D depicts examples of low (left) and high (right) entropy. On average, protein levels of most proteins were greater in cancer compared to immune and stroma cells (Figure 2) which allows more states and leads naturally to high entropy in cancer cells.

**Figure 7.**
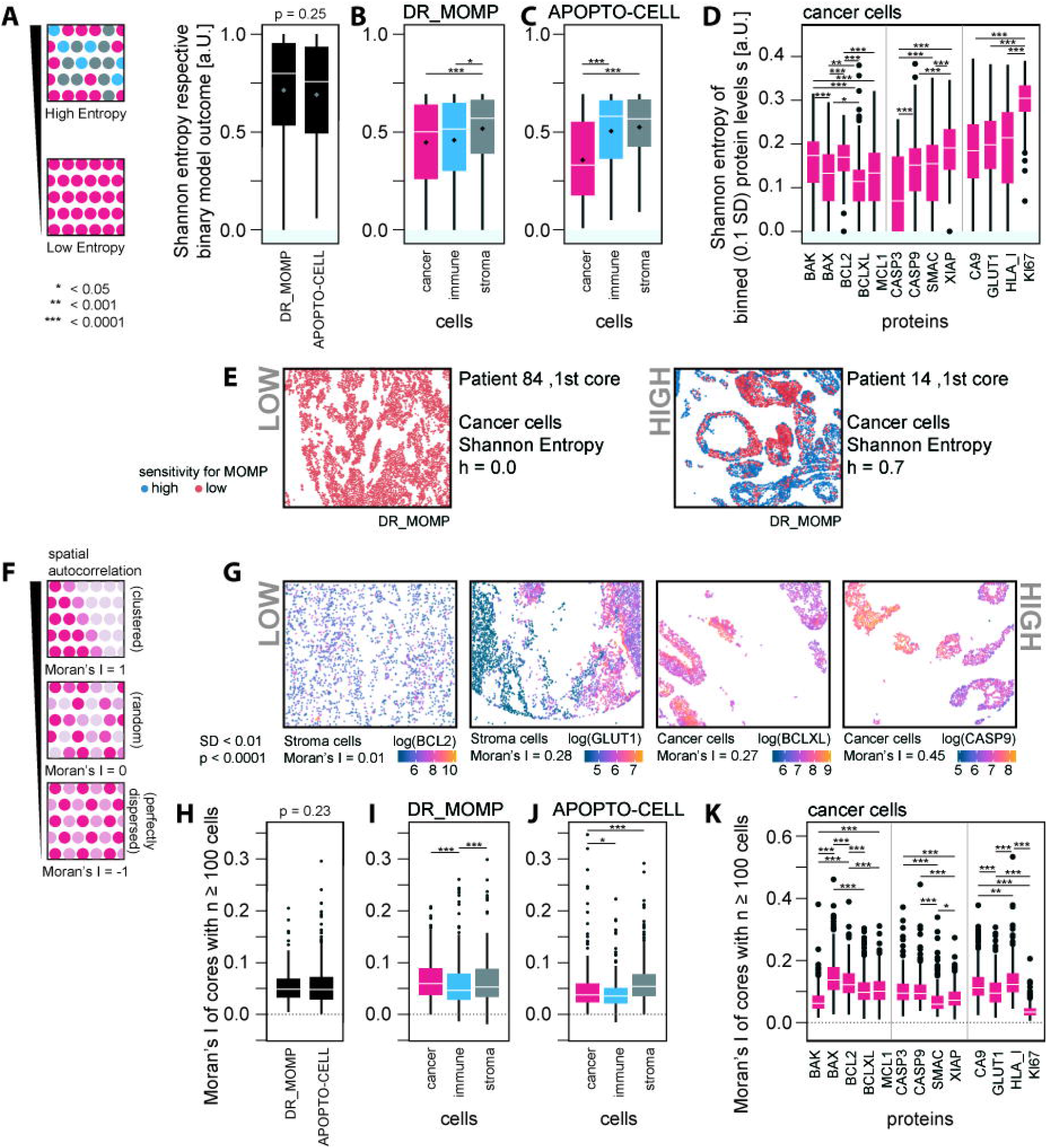
Heterogeneity analysis calculating cells’ (A-E) Entropy and (F-K) Moran’s I for apoptosis model predictions as well protein levels. (A-E) Entropy (information theory) is a measurement for the bias to one state, (A) with low entropy marking captaincy for a one state and high entropy marking uncertainty for one or multiple states. We first determined the binary Shannon entropy for (B) low/high sensitivity for MOMP (DR_MOMP) and (C) low/high caspase activity (APOPTO-CELL; ANOVA, Tukey post-hoc), finding surprisingly significant lower entropy in cancer cells (red) compared to immune (blue) and stroma cells (gray). (D) Subsequently, we calculated the Shannon Entropy for the proteins using bins for protein level with a bin width of z-score = 0.1 SD for each protein respectively. The calculated Shannon Entropy for stroma and Immune cells can be found in supplementary figure 5. (E) shows examples with low (left) and high (right) entropy for the DR_MOMP model. (F) Moran’s I is a measurement of spatial autocorrelation with a Moran’s I approaching 0 and < 0 indicating spatial dispersion and a Moran’s I approaching 1 marking spatial clustering. Panel G shows examples of protein levels with low (left) and high (right) Moran’s Is. (G-K) We determined cores’ Moran’s I for low/high (I) sensitivity for MOMP, (J) caspase activity and (K) respective protein levels (in cancer cells). Calculated Moran’s I for Stroma and Immune cells can be found in supplementary figure 6. (G) While a Moran’s I around 0 shows no spatial autocorrelation, values around 0.2 or greater indicate presence of local spatial autocorrelation within the cores.

We also assessed the presence of systematic spatial variation of protein levels and model predictions (spatial autocorrelation) by measuring Moran’s I in each core (Figure 7F-J). A Moran’s I of 1 indicates a perfect spatial separation (e.g. left *versus* right separation), while a value of −1 indicates a perfect dispersion (checkerboard pattern; Figure 7F). A Moran’s I is close to zero for a random distribution. Figure 6G depicts examples of low (left) and high (right) Moran’s I.

Overall, we found little evidence of strong spatial autocorrelation or spatial separation in the majority of cores suggesting that cells that were close to each other did not have similar protein levels or similar apoptosis sensitivities. Overall, we did not find any statistically significant difference in Moran’s I between the different apoptosis models (Figure 7H). Similar to the Entropy, this value is biased if cells of different types are spatial separated, and Moran’s I needs to be studied individually. Although we observe only minor difference for the DR_MOMP model (Figure 7I), cancer cells with different predictions for APOPTO-CELL were significantly more randomly dispersed compared to stromal cells (ANOVA p = 0.004 and Tukey post-hoc p = 0.002; Figure 7E).

Calculating Moran’s I for cells’ protein levels, we found that the majority of cancer cells have a score less than 0.2 suggesting a tendency towards a low correlation between protein level and the distance between cells (Figure 7K). However, individual cores showed high spatial clustering for individual protein suggesting that neighboring cells are more likely to have similar protein levels than distant cells in these cores. Among the proteins relevant for DR_MOMP, BAX and BCL2 showed the higher Moran’s I compared to BAX, BCL(X)L and MCL1 (Figure 7K). Among proteins used in the APOPTO-CELL model, SMAC had the lowest Moran’s I compared to PRO-CASPASE 3, 9 and XIAP (Figure 7K). Of note, since immune cells are more mobile than epithelial or stroma cells, we would assume to find the lowest Moran’s I in these cells. However, this was only the case for BAK, BCL2, PRO-CASPASE 3 and GLUT1 (Suppl. Figure 6). While numerically different, overall the Moran’s I was similar for BAX, BCL(X)L, MCL1, SMAC and CA9 if stratified for cell types. We observed the greatest difference between cells of different types for BAK, BCL2, GLUT1, HLA-I and KI67 (Suppl. Figure 6).

Collectively, intra-tumoural heterogeneity in apoptosis signaling was surprisingly not increased in cancer cells when compared to leukocytes and other stromal cells, suggesting that heterogeneity in apoptosis signaling represents an intrinsic, non-genomic cell property that is not increased by the process of malignant transformation.

## Discussion

The present study constitutes the first report describing the quantitative and spatial distribution of key mitochondrial apoptosis proteins at single cell resolution in intact cancer tissue. Using multiplexed immunofluorescence imaging (MxIF) we provide information on 2.4 million apoptosis protein profiles in six different cell types and deliver the first atlas of apoptosis signaling proteins in a large cohort of patients (164 colorectal cancer patients). We furthermore conducted a systems-based analysis of each individual cell’s apoptosis sensitivity. Our dynamic systems modelling estimated that cancer cells were generally more sensitive to apoptosis signaling than immune or stromal cells, however with significant heterogeneity between patients. We also characterized the level of intra-tumoral heterogeneity in apoptosis signaling in colorectal cancer, and demonstrate that intra-tumoral heterogeneity in apoptosis signaling was not increased in cancer cells when compared to leukocytes and other stromal cells.

### Apoptosis protein mapping in colorectal cancer and its implication for future therapy

Our first analysis steps constituted the mapping of protein profiles to the different cell types present in the tumor microenvironment. Multiplexed protein imaging has been increasingly used as a tool for spatial analysis of tumor cell types and microenvironment over the last 10 years (Angelo et al., 2014; Gerdes et al., 2013; Goltsev et al., 2018; Gut et al., 2018; Kalra and Baker, 2017; Rashid et al., 2019; Saka et al., 2019; Tan et al., 2020) and there are increasing number of multiplexing methods for in situ RNA and DNA detection (Decalf et al., 2019; Kishi et al., 2019; Moffitt and Zhuang, 2016), Cell DIVE has been used to analyze tumor cell heterogeneity in CRC (Badve et al., 2021; Spagnolo et al., 2017), ductal carcinoma *in situ* (DCIS) (Badve et al., 2021; Gerdes et al., 2018), breast cancer (Sood et al., 2016), glioma and glioblastoma (Berens et al., 2019) and melanoma (Yan et al., 2019). Unlike standard immunohistochemistry methods which are limited to 1-5 markers in a single section, multiplexed immunofluorescence imaging methods can provide single cell data on up to 60 proteins in a single sample, including cell spatial coordinates, thus allowing analysis of co-expressed biomarkers and relationships between cells types and functional status, as described in this paper.

Overall, we found that the ‘average protein level’ in a core (of the proteins we investigated), as evaluated in bulk assays, is predominantly due to by the signal from cancer cells compared to immune or stroma cells (Figure 2BE). However, among the analyzed key proteins regulating mitochondrial apoptosis, we found interesting differences between the cell types (Figure 2AD). One of the key findings was an enrichment in BCL-2 protein levels in immune cells when compared to cancer and other stromal cells. This finding may have important implications regarding the use of BCL2 antagonists such as Venetoclax for the treatment of solid tumors. Venetoclax is well tolerated in patients with relatively few side effects and would represent an ideal adjuvant and sensitizer to chemotherapy for the treatment of chemotherapy-resistant solid tumors.

Nevertheless, Rohner *et al. (Rohner et al., 2020)* have previously shown that inhibition of BCL2 by ABT-199 caused cell death in all types of lymphocytes but specifically reduced the counts of B cells in humans. In addition, the authors showed that while T cells showed equivalent high levels of BCL2, latter were significantly less affected compared to B cells, emphasizing that triggering apoptosis is “the sum of the interplay of a network of anti-and pro-apoptotic BCL-2 family members” (Rohner et al., 2020) and highlighting the importance of a systematical assessment of apoptosis. Our data suggest that in certain patients BCL2 is also highly expressed in epithelial cancer cells, which could suggest that BCL2 antagonist therapy may effectively reduce the overall anti-apoptotic threshold of cancer cells. Due to complexity in cell-type specific BCL2 expression, our study suggests that evaluation of BCL2 levels in bulk tissues samples may not be sufficient as a stratification tools for BCL2 antagonists.

We found that MCL1 levels were enriched in epithelial and immune cells, with significantly lower levels in stroma tissue. As MCL1 antagonists are also being currently developed as apoptosis sensitizers for MCL-1-dependent cells, effects of MCL1 antagonists on immune cells may also need to be considered. Of note, quantitatively, MCL1 levels were lower in cancer cells when compared to the anti-apoptotic proteins BCL2 and BCL(X)L. Another interesting aspect of our study was the strong enrichment of BAK in cancer cells. Recently, agents have been developed that activate BAX and BAK directly (Walensky and Gavathiotis, 2011), including molecules that do not interact with the BH3-binding pocket of anti-apoptotic proteins or pro-apoptotic BAK and induces cell death in a BAX-dependent fashion (Gavathiotis et al., 2012; Gavathiotis et al., 2008). Our results suggest that BAK in particular is a good target in colorectal cancer. Cancer cells also had significant higher levels of SMAC, XIAP and PRO-CASPASE 9 compared to immune or stroma cells. It is therefore possible that colorectal tumors expressing high XIAP levels in cancer cells are effectively sensitized by SMAC mimetics (Fichtner et al., 2020), however analysis of both XIAP and SMAC levels may be required for future patient stratification.

### Priming of cancer cells and the degree of inter-individual heterogeneity

We also utilized the protein profiles for calculations of apoptosis sensitivity at the systems level. Because of the complexity of apoptosis signaling with multiple signaling redundancies and feed-back signaling, several groups have developed functional or computational models that describe apoptosis sensitivity on a systems level. One such as approach, termed ‘*BH3-profiling’*, interrogates the response of the mitochondrial apoptosis pathway to pro-apoptotic BH3-only protein peptide mimetics (Certo et al., 2006; Del Gaizo Moore and Letai, 2013; Montero and Letai, 2018). However, this technique requires fresh tissue and living cells. To enable analysis of fresh frozen or formalin-fixed paraffin-embedded tissue, we developed DR_MOMP (Lindner et al., 2013) as an ODE-based model of MOMP that, similarly to BH3 profiling, calculates the response of the BCL2 signaling network to BH3-only proteins activated upon cellular stress (Flanagan et al., 2015; Lindner et al., 2013; Lindner et al., 2017). It has been extensively validated experimentally in colon and other cancer cells (Lindner et al., 2013; Lindner et al., 2017; Lucantoni et al., 2018). Furthermore, we developed APOPTO-CELL as an ODE model that calculates the sensitivity of cells to activate caspase-3 downstream of MOMP (Huber et al., 2007; Rehm et al., 2006), as this process represents an important second control step. APOPTO-CELL has also been extensively validated in-house using single cell imaging and population-based approaches in cervical, colorectal and glioblastoma cells (Murphy et al., 2013; Salvucci et al., 2019a; Salvucci et al., 2017; Schmid et al., 2012). Both models have also been shown to predict responses to apoptosis sensitizers in preclinical settings (Lucantoni et al., 2018; O’Farrell et al., 2020). Our study supports the previously developed concept that tumor cells are indeed ‘primed’ to undergo mitochondrial apoptosis (Llambi et al., 2011; Ni Chonghaile et al., 2011). Including additional markers for BH3-only proteins and caspase-independent cell death pathways will allow us to gain a more holistic picture of possible cell fates in the future.

However, a surprising observation was that (based on model prediction) this appeared to result predominantly from an enhanced ability to overcome both apoptosis barriers, MOMP and activation of caspase-3 activation downstream of MOMP, as immune cells lack sensitivity for MOMP and other stromal cells showed less sensitivity to caspase-3 activation (Figure 6AC). When comparing the ability to undergo MOMP, cancerous cells were equally sensitive to stromal cells to undergo MOMP. In contrast, immune cells appeared to be highly resistant to MOMP due to their relatively high expression of increase in BCL2. Interestingly, we found that cytotoxic (CD8+) T cells were overall significantly more susceptible to apoptosis stimuli compared to other immune cells. This is clinical relevant since tumor infiltration by cytotoxic T Cells has been found to be significantly positively correlated with better survival in colorectal cancer(Naito et al., 1998). In patients with breast cancer, changes in the ratio between FOXP3+ (Regulatory) and CD8+ (cytotoxic) T cells (before and) after neoadjuvant chemotherapy was highly associated with clinical response (Ladoire et al., 2008). Similarly, a low density of cytotoxic T Cells in tumor tissue after chemotherapy was associated with poor response in patients with rectal cancer (Matsutani et al., 2018). Therefore, increased risk of apoptosis of cytotoxic T cells (e.g. following chemotherapy) may abrogate these benefits.

Nevertheless, our study cannot address the questions whether cancer cells are capable of activating more BH3-only proteins at a given genotoxic (or metabolic) stress dose. We also observed significant patient-to-patient heterogeneity in apoptosis sensitivity at both levels, while core-to-core differences within single patients were less pronounced. Moreover, our combined analysis of both apoptosis signaling pathways in each individual cell also allowed us to investigate potential blocks in either of these pathways. Our combined analysis showed that the majority of cancer cells showed a high sensitivity for at least one of the two apoptosis pathways up-and downstream of MOMP, which was not observed to a similar degree in immune or other stromal cells.

### Intra-tumoral heterogeneity

One of the limitations of the current study was that tumor core regions were analyzed, while tumor margins in the invasive zone were not investigated. However other studies have pointed to the importance of core regions in tumor progression due to silencing/methylation as a consequence of tissue hypoxia (Thienpont et al., 2016). Based on this and other previous studies pointing to an importance of intra-tumor heterogeneity in tumor progression and resistance, we also investigated intra-tumor heterogeneity in apoptosis signaling. Collectively, our entropy and spatial image analyses of the mitochondrial apoptosis pathway did not suggest that cancer cells showed an increased cell-to-cell or spatial heterogeneity when compared to immune or other stromal cells. However, as shown in the examples for Moran’s I (Figure 7G), there can be a significant different between the value of 0.0 and 0.2 and assessing autocorrelation with alternative methods, such as Variograms, may be of benefit. Another limitation was that we resolved the cell’s phenotype in only three classes. Observed heterogeneity in predicted model response and measured protein levels could arise through a high number of various differentiated cells, and cells of the same type might have significantly lower heterogeneity if compared among each other. Notwithstanding these limitations, our studies indicate that intra-tumoral heterogeneity in apoptosis signaling was not increased in cancer cells, suggesting that this represents an intrinsic, non-genomic property not increased by the process of malignant transformation. This observation is supported by earlier studies in cell lines which demonstrated the importance of non-genomic heterogeneity in apoptosis signaling due to fluctuations in protein levels over the lifetime of a cell. Rehm *et al. (Rehm et al., 2009)* reported that sibling cells underwent apoptosis execution within a narrow time window and that random cell pairs were significantly less synchronous in undergoing apoptosis, independent of activating the intrinsic or extrinsic pathway. However, the authors also reported that neither cell-to-cell distance nor cell membrane contacts influenced the synchrony in apoptosis execution of sibling cells (Rehm et al., 2009). Similarly, Spencer *et al. (Spencer et al., 2009)* previously showed that differences in the protein levels regulating apoptosis are the primary causes of cell-to-cell variability in probability of death, with the protein state being transmitted from mother to daughter, and protein synthesis rapidly promoting divergence between these cells.

While we here consider the levels of 9 apoptosis markers, we did not take into account proteins’ state such as BCL2’s phosphorylation status (Ruvolo et al., 2001) nor subcellular localization of proteins which is possible to account for with the Cell DIVE™ platform. For example, BAX’s localization at the mitochondria or in the cytosol was reported to be clinically relevant in acute myeloid leukemia (Reichenbach et al., 2017) and hepatocellular carcinoma (Funk et al., 2020). BAX localization could be considered by including a mitochondrial marker, or by analyzing the BAX signal within the cytosolic cell mask, with an evenly distributed signal suggesting cytosolic localization, and uneven distribution suggesting localization at mitochondria.

In conclusion, our study provides the first map of apoptosis sensitivity at individual protein and systems level in intact colorectal cancer tissue. We holistically describe both patient-to-patient and intra-tumor heterogeneity in apoptosis signaling in stroma, immune and cancer cells which has important implications for the future use of apoptosis sensitizers in the treatment of colorectal cancer.

## Acknowledgments

This work was funded by a US-Northern Ireland-Ireland Tripartite grant from Science Foundation Ireland and the Health Research Board to JHMP (16/US/3301) and the National Cancer Institute (Systems Modeling of Tumor Heterogeneity and Therapy Response in Colorectal Cancer; to FG). EPO’C is support by an RCSI Bon Secours Hospital MD StAR fellowship and the Beaumont Hospital Cancer Research and Development Trust. DBL, PD, ML, XS were supported by US-Ireland R01 award (NI Partner supported by HSCNI, STL/5715/15).

## Author Contributions

AUL, MS, EMcD, SaCh, StCa, DPO’C, ADC, ASP, PLP, ML, AS, AS and JFG were involved in methodology, data validation and curation. EMcD, SaCh, ADC, ASP, AS, JFG and FG performed the Cell DIVE processing as well as cell segmentation and single cell quantification. AUL and MS statistically analyzed the data and study investigation. JPB, DAmcN, SVS, JHMP, PLP, PD and DBL were involved in clinical sample acquisition and data collection. EMcD AC and ASP conducted the sample imaging and image processing. ASP and AS led the single cell analysis workflows for epithelial and immune cell analysis. StCa and MF grow and processed the control cell lines. AUL, MS, JFG, SaCh, DBL, XS, FG and JHMP reviewed the data. AUL, MS, MR, DBL, FG and JHMP designed experiments .AUL and JHMP wrote the manuscript. AUL created the manuscript figures. MR, DBL, FG and JHMP were involved in funding acquisition. All Authors edited and revised the manuscript text.

## STAR★Methods

### Key Resource Table

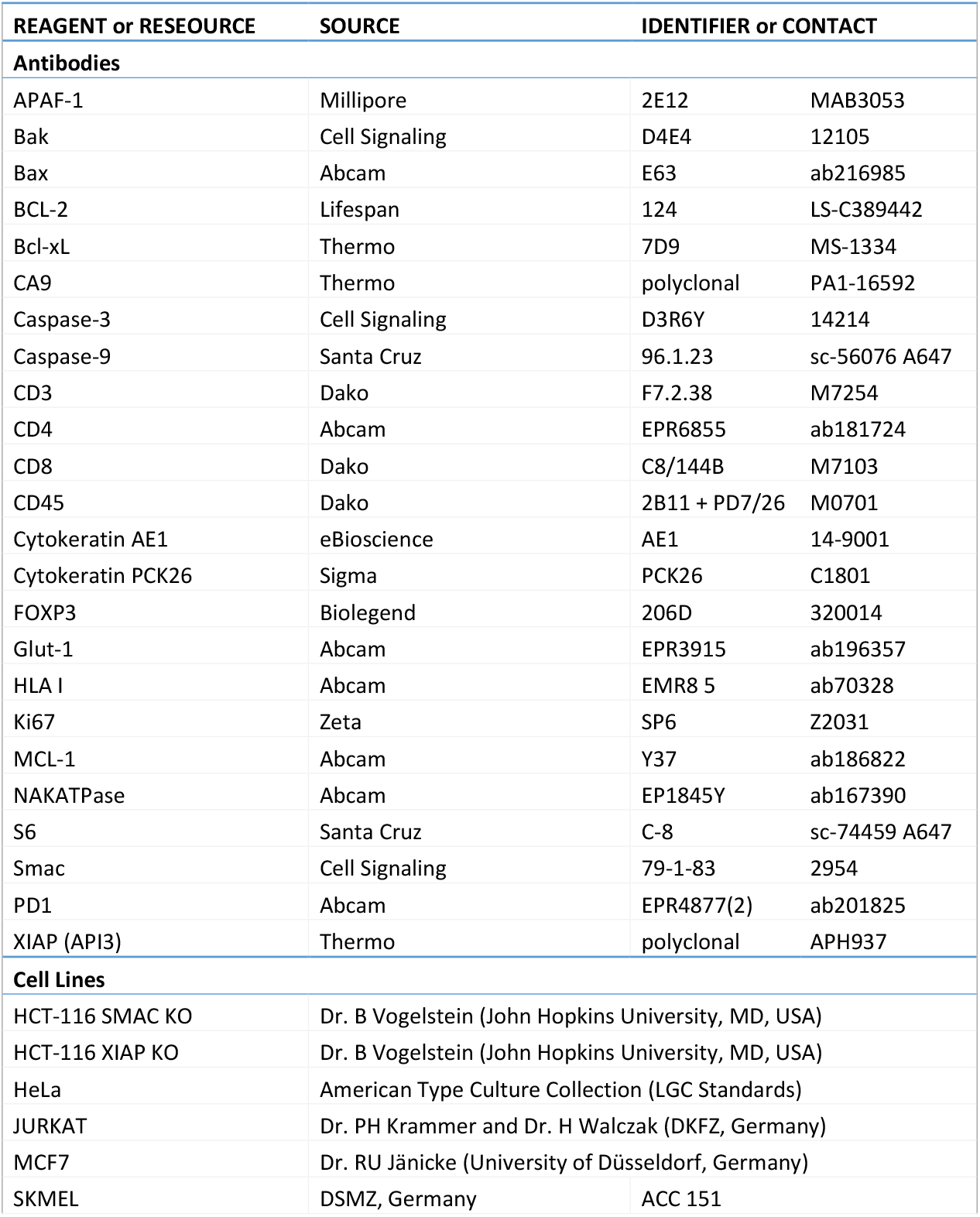
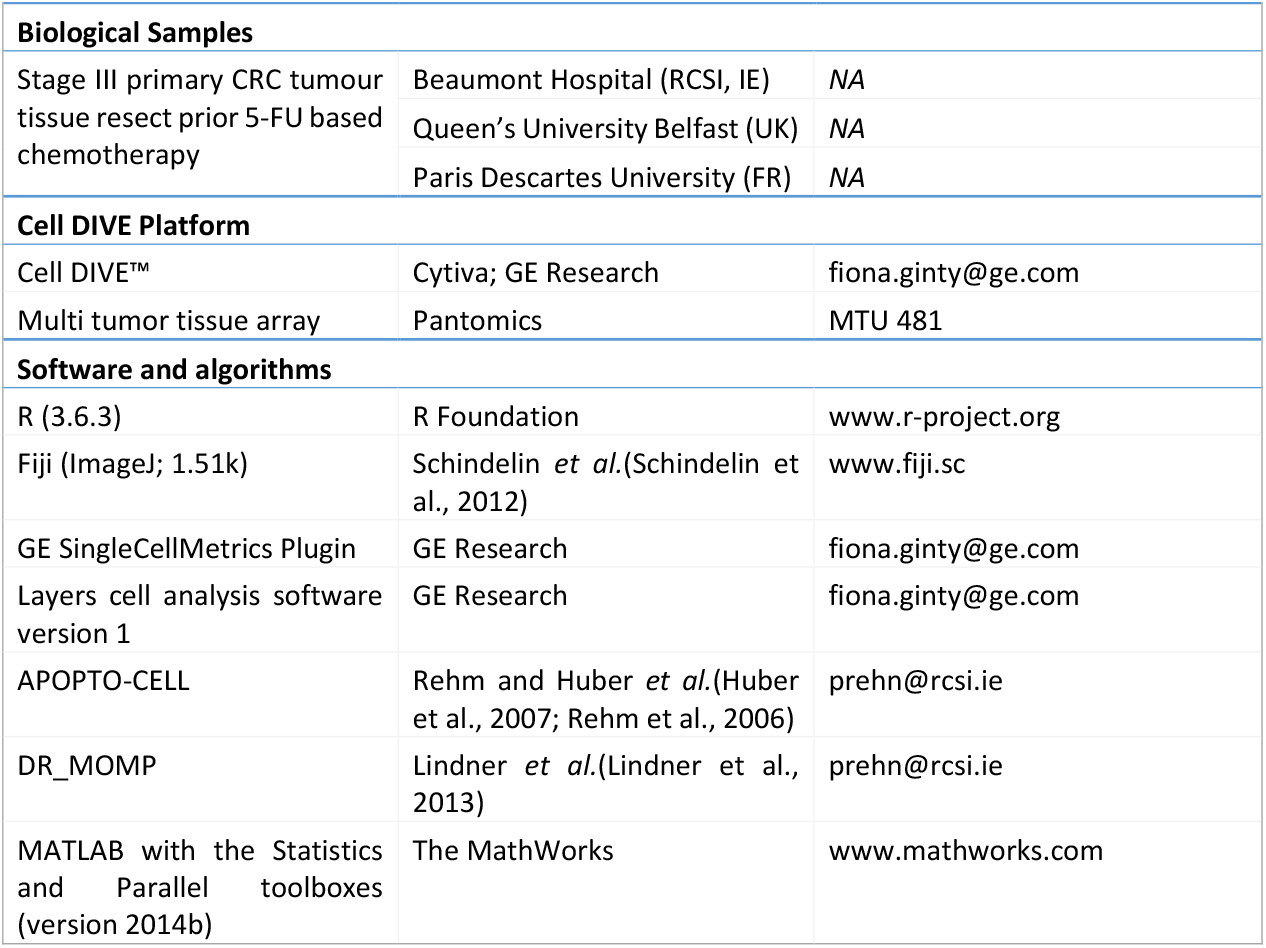

#### Resource Availability

##### Lead Contact

Further information and request for code or resources should be directed to and will be fulfilled by the lead contact, Prof. Jochen Prehn.

##### Materials Availability

- This study did not generate new unique reagents.

##### Data and Code Availability

- Imaging data, cell masks and generated single cell measurements of 20 markers is available from the lead contact.
- The full pipeline for data analysis is available from the lead contact.
- Any additional information required to reproduce this work is available from the Lead Contact.

#### Experimental Model and Subject Details

This section does not apply to our computational study.

## Method Details

### Colorectal cancer cohort

Formalin-fixed, paraffin-embedded (FFPE) primary tumor tissue sections were obtained from 170 chemotherapy-naïve, resected stage III CRC patients. Tumor samples were collected from three centers: Beaumont Hospital (RCSI, Ireland), Queen’s University Belfast (UK) and Paris Descartes University (France). All centers provided ethical approval for this study and informed consent was obtained from all participants. A summary of the clinical characteristics of the cohort is provided in Suppl. Table 1. Data of 46 cores of 36 patients were dropped after quality assessment of the stained tissue (see below). All cores of two patients were removed in this process.

### Cell lines

Three technical replicates (cores) of pellets of formalin-fixed HeLa, Jurkat, MCF7, SKMEL, HCT-116 SMAC^KO^ and HCT-116 XIAP^KO^ cells in which quantities of mitochondrial apoptosis proteins were previously determined (Lindner et al., 2013; Passante et al., 2013; Rehm et al., 2006) were included in the construction of the tissue microarray (TMA) in parallel to the patients’ cores, and served as quality control and internal standards for protein quantification. 3 of 18 cores of two cell lines were removed after quality control. Cells were grown to 80% confluence. Media was replaced 12-24 hours before fixation. To fix cells, cells were gently washed in sterile 1XPBS solution. Cell monolayers were covered with 5 mL 10% neutral-buffered formalin (NBF) for 2-5 min. Cells were scraped into NBF, and collected into labelled 50 mL tubes, and stored at 4 C for at least 3-4 hours. For further processing, cells were centrifuged at 1,200 rpm for 5 min and washed in 1% low melt agarose solution XBPS before re-suspension in 0.5 ml 80% ethanol and centrifugation at 12,000 rpm twice for 5 min. Subsequently 80% of ethanol was aspirated and cell pellets were molded into caps and frozen, prior to TMA construction.

### Antibody validation and conjugation

Commercially acquired antibodies underwent multi-step process of validation and conjugation (as previously described by Gerdes *et al.* (Gerdes et al., 2013). Briefly, at least 2-3 clones for each target were stained in parallel using a multi-tissue array (MTU 481, Pantomics, CA) and staining performance visually compared. At least one antibody clone was down-selected for conjugation with either Cy3 or Cy5 bis-NHS-ester dyes. Epitopes were also tested for sensitivity to the dye inactivation solution (basic hydrogen peroxide) by exposing multi-tissue arrays to 0, 1 and 10 rounds the solution and stained with the antibody of interest and compared. Approx. 10% of epitopes have been shown to have decreased signal following exposure to the inactivation solution and those antibodies are placed early in the multiplexing sequence (Gerdes et al., 2013). The key resource table shows the antibodies, clones and conjugates used in this study. Briefly the markers and staining rounds were as follows: Round 1: BCL2, APAF1; Round 2: MCL1, PRO-CASPASE-9; Round 3: S6, PRO-CASPASE-3; Round 4: BAX, SMAC; Round 5: BAK, XIAP; Round 6: NaKATPase, BCL(X)L; Round 7: Cytokeratin PCK26, CD8; Round 8: Cytokeratin AE1, FOXP3; Round 9: CD4, Ki67; Round 10: HLA1, CD45; Round 11: Glut1, CA9; Round 12: CD3, PD1; Round 13: S6 (repeated). Note that in total, 9 background imaging rounds were also included.

### Immunofluorescence Imaging of Patient TMAs

Multiplexed immunofluorescence iterative staining of the CRC TMAs was performed as previously described (Gerdes et al., 2013) using the Cell DIVE™ technology (Cytiva, Issaquah, WA; formerly GE Healthcare). This involves iterative staining and imaging of the same tissue section with 60+ antibodies and is achieved by mild dye oxidation between successive staining and imaging rounds. In total, there were 13 staining rounds using the antibodies described above and DAPI was imaged in each round. The Leica Bond (Leica Biosystems) was used for antibody staining and the IN Cell 2200 was used for imaging. Staining and image recording was repeated twice for S6 due to staining failure. Exposure times were set to fixed values for all images of a given marker.

### Image pre-processing

Immunofluorescent images were processed and cells were segmented and quantified as described previously (Gerdes et al., 2013). To summarize, cells in the epithelial and stromal compartments were segmented using DAPI, pan-cytokeratin, S6, and NaKATPase stains (Gerdes et al., 2013). Images and segmented cell data then underwent a multistep review process (described by Berens et al.(Berens et al., 2019): 1) images were visually reviewed and manual scoring of tissue quality and segmentation was determined by at least one researcher. Images with poor quality staining or too few cells were excluded from data analysis; 2) cell filtering based on minimum and maximum number of pixels in each sub-cellular compartment (> 10 pixels and < 1500 pixels per compartment) and 1-2 nuclei per cell; cells with values outside these limits were removed 3) confirmation of excellent alignment of all cells in all staining rounds compared to the first round of staining. For this, an automated QC score was generated for every cell in each imaging round by correlating baseline DAPI images with all corresponding DAPI images from other multiplexing rounds. A perfect score of 1 indicated perfect registration, no cell loss and no cell movement. A score of 0 indicated complete loss of that cell after baseline imaging. After quality control, cells included in the analysis had a median QC score of 0.95, with 53% having a QC score greater than 0.8. The average QC score was 0.57. In comparison, 83% of cells removed during quality control had a QC score less than 0.1 with an average QC score of 0.15. From the single-cell segmentation masks, the mean intensity, standard deviation, and coherent statistics were quantified for each protein with respect to the whole cell as well as xy-location. From the single-cell segmentation masks, the mean, standard deviation, median, and maximum staining intensity for each protein were quantified with respect to the whole cell, cell membrane, cytoplasm, and nucleus as well as cell location, area, and shape. Following quantification, slides were normalized for batch effects and exposure time for each channel/marker analyzed. 48 positions showing major cell loss during staining rounds were excluded from all analysis, as well as cells within the images’ margins of 15 pixel on the x-axis and 10 pixel on the y-axes were dropped from all data analysis. 74 positions showing major or minor cell loss during staining rounds were excluded from training datasets for post-processing such as batch correction or cell classification.

### Post pre-processing and batch correction

To correct for a possible batch effects between slides, cells’ mean intensity were first normalized using upper-quantile normalization, grouped by protein marker and slide. Secondly, quantiles of the normalized intensities were plotted against their rankits, and an affine transformation matrices to rotate the function to the main diagonal were calculated. Obtained transformation matrices were applied on the intensities, and pixel intensity values were restored using linear regression and upper-quantile normalized values. Solely for the batch correction, cells within 5% of the images’ margins were excluded for the calculation of the reference values. The batch correction was quality controlled with cell lines spotted in parallel to tissue samples on 3 of 5 slides.

### Immune Cell classification

To differentiate cell types, we used CD3, CD4, CD8, CD45, FOXP3, PCK26 and Cytokeratin AE1 markers. We manually annotated 4,839 AE1-or PCK27-positive cells as (epithelial) cancer cells. Of 3,121 CD3-positive cells (Beare et al., 2008), 788 CD4-positive cells were annotated as Helper T cells (Beare et al., 2008), 991 CD8-positive cells were annotated as Cytotoxic T cells (Beare et al., 2008), and 1,360 FOXP3-positive cells were annotated as Regulatory T cells (Hori et al., 2003). 3,369 CD3-negative cells that were either CD4-, CD45-or CD8-positive were annotated as other leukocytes. 3,837 cells that lacked any marker but were DAPI positive were annotated as stroma-rich cells (other stromal cells). Using the manual annotations, we constructed a random forest of 2,000 trees (R package *randomForest*, version 4.6-14) and employed it to classify all cells.

### Protein profiling and apoptosis sensitivity modelling

Protein levels of BAK, BAX, BCL2, BCL(X)L and MCL1 were normalized to the mean protein levels in HeLa cells spotted in parallel to patients’ core on 3 of 5 slides. Protein’s molar concentrations were calculated using previously established HeLa concentrations (Lindner et al., 2013).The five proteins were used as input for the DR_MOMP mathematical model(Lindner et al., 2013) that models the BCL2 signaling pathway before MOMP and is able to calculate the stress dose required for MOMP or if a cell undergoes MOMP due to a specified stress. DR_MOMP (Lindner et al., 2013) was translated from its MATLAB implementation to C++ and R using deSolve (1.28), doParallel (1.0.15) and Rcpp (1.0.5).

APOPTO-CELL (Rehm et al., 2006) was executed in MATLAB with the *Statistics and Parallel toolboxes* (version 2014b, The MathWorks, Inc., Natick, MA, USA). The model requires molar concentrations [µM] of APAF1, PRO-CASPASE 3, PRO-CASPASE 9, SMAC and XIAP as input to predict amount of cleaved substrate, as a readout for apoptosis susceptibility [%]. Previous research(Hector et al., 2012; Salvucci et al., 2019a) has shown that APAF1 is not the limiting factor in apoptosome formation in the CRC settings(Hector et al., 2012; Salvucci et al., 2019a) and was set to 0.123 µM. Molar protein concentrations for PRO-CASPASE 3, PRO-CASPASE 9, SMAC and XIAP were estimated by aligning signal intensities [a.U.] to profiles [µM] determined in a reference clinically-relevant CRC cohort(Hector et al., 2012) with an established pipeline(Salvucci et al., 2019a; Salvucci et al., 2017). The pipeline was built upon the assumptions that 1) measurement ranking is preserved (monotonic relationship between batch-corrected signaling intensities and molar concentrations); and 2) absolute concentration profiles in clinically-matched cohorts are comparable. The pipeline implementation follows directly from the above assumptions. Briefly, for each protein smoothed kernel probability distribution objects were fitted to 1) the known protein molar concentrations of the reference CRC cohort (Hector et al., 2012) and 2) batch-corrected multiplexed signal intensities (restricted to high quality data points where no signal loss across staining rounds had been observed), with the MATLAB function *fitdist (*as detailed in Salvucci M *et al. (Salvucci et al., 2017)*). The inverse cumulative distribution transformation of the reference distribution kernel was applied on the batch-corrected signal intensities to determine the corresponding absolute concentrations (MATLAB function *icdf*).

For both models, we performed two sets of simulations: 1) per-core and 2) per-cell. For the per-core simulations, we aggregated (by median) the batch-corrected protein intensities across all cells for each core per patient prior to conversion to molar concentrations, resulting in one simulations per-core and thus 2-3 simulations per patient. For the per-cell simulations, we performed a simulation for each cell, totaling ∼3.5 million simulations for 164 patients included in the study.

### Statistical Analysis

All statistical tests were performed in R (3.6.3) and p values of < 0.05 were considered statistically significant. All data are presented as mean ± SEM. All statistical tests were performed in R. If not otherwise mentioned, two-tailed t tests were performed for pairwise comparison, while analysis of variance (ANOVA) with Tukey honest significance post-hoc tests were performed in cases of the comparison of three or more populations. The quartile coefficients of dispersion (COF) were calculated using (Q_3_ -Q_1_) / (Q_3_ + Q_1_) with Q_n_ be the respective quartiles. Shannon Entropy was calculated either using log_2_ for binary populations or the natural logarithm, with 10^−10^ added to all values. Moran’s I was calculated using the R package *ape* (5.4-1) without outliers and only on populations > 100 cells. Distances > 2,000 px were set to 2,000 px. Consensus Clustering was performed using *ConsensusClusterPlus* (1.48.0) with a seed of 42, 100,000 repetitions, Spearman and Ward’s method as parameters. For the bootstrap analysis, slides were randomly 100,000 times randomly paired using a seed of 42.

## Supplemental Information titles and legends

Supplementary Table 1 – Patient information with mean cell fractions and DR_MOMP and APOPTO-CELL results for aggregated protein levels for patient-matched cores.

Supplementary Table 2 -of transcriptional data derived from flow-sorted immune, epithelial and fibroblast populations isolated from CRC primary tumor tissue (GSE39396).

Supplementary Figure 1 -Plot of patients’ consensus cluster score of patient-matched cores after hierarchical consensus clustering using cancer, immune and stroma cell fractions of each core. Patients with a low consensus score (0) show high difference in cell fractions between matched cores while patients with a high consensus score (1) show high similarity in cell fractions between matched cores.

Supplementary Figure 2 – Box plot of transcriptional data derived from flow-sorted immune (n = 6), epithelial (n = 6) and fibroblast (n = 6) populations isolated from CRC primary tumor tissue (GSE39396 (Calon et al., 2012); Suppl. Table 2; ANOVA and Tukey post-hoc).

Supplementary Figure 3 -Box plot of quartile coefficients of dispersion of protein levels of each core and stratified for cancer (red), immune (blue) and stroma cells (grey).

Supplementary Figure 4 -In analog to the correlation plot in Figure 4C-E showing the median correlation coefficient in all (black), cancer (red), immune (blue) and stroma (gray) cells, but including the interquartile range.

Supplementary Figure 5 -Calculated the Shannon Entropy for the proteins using bins for protein level with a bin width of z-score = 0.1 SD for each protein respectively and stratified for cancer (red), immune (blue) and stroma (gray) cells. Proteins were sorted base for (A) DR_MOMP, (B) APOPTO-CELL and (C) others.

Supplementary Figure 6 -Calculated cores’ Moran’s I for low/high (I) sensitivity for MOMP for protein levels stratified for cancer (red), immune (blue) and stroma (gray) cells. Proteins were sorted base for (A) DR_MOMP, (B) APOPTO-CELL and (C) others.

