## Supplementary figures and images for "An atlas of inter- and intra-tumor heterogeneity of apoptosis competency in colorectal cancer tissue at single cell resolution"

### Supplementary Figure 1

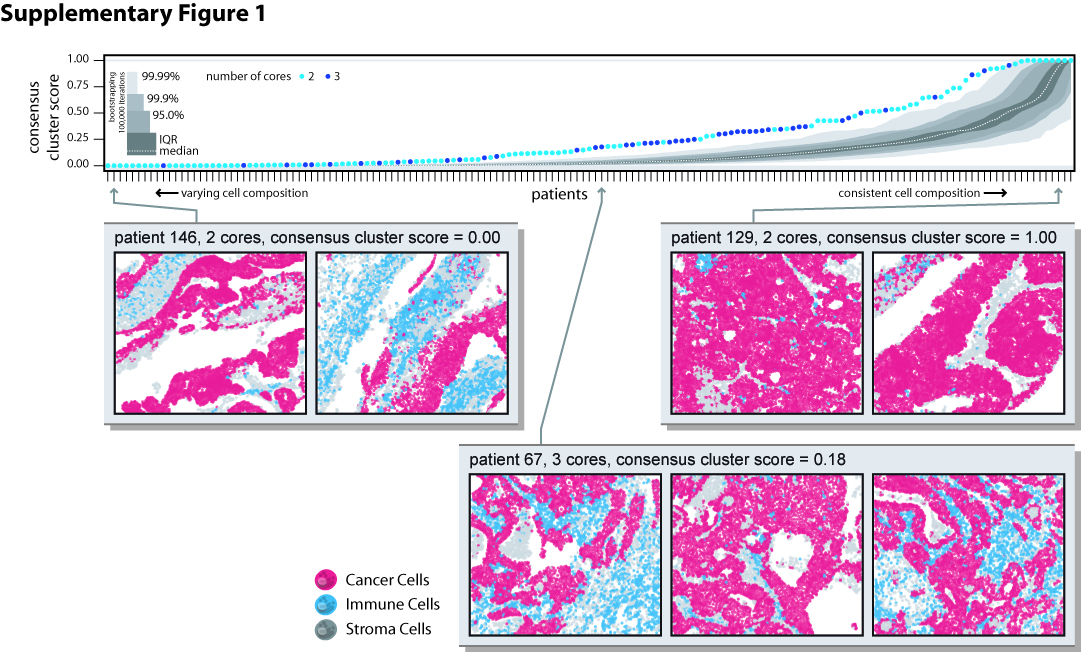

### Supplementary Figure 2

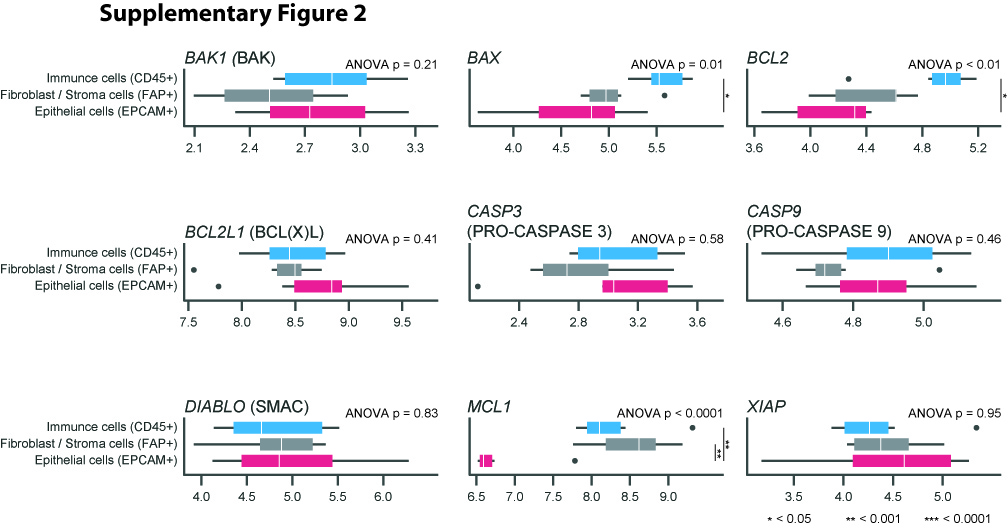

### Supplementary Figure 3

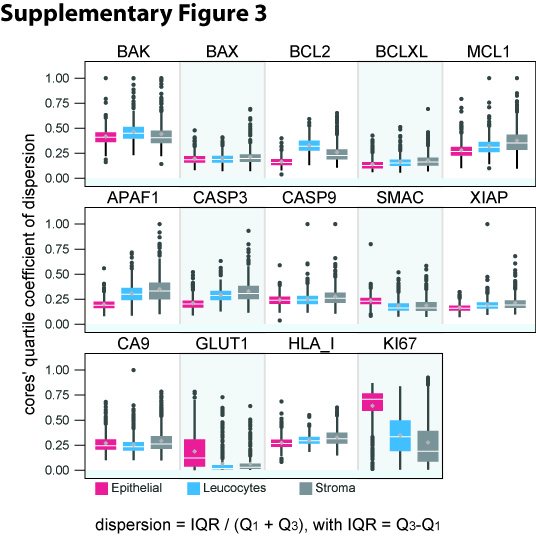

### Supplementary Figure 4

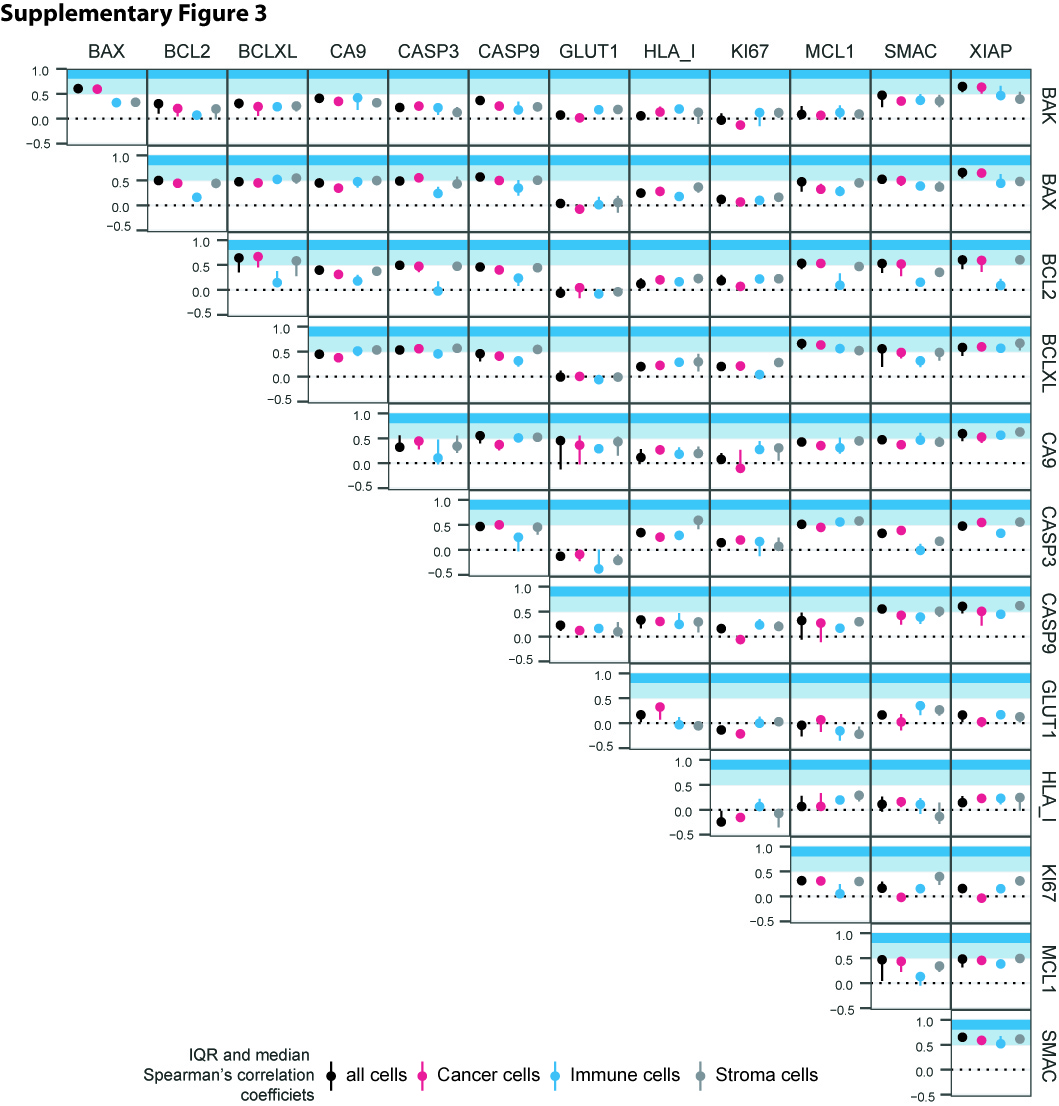

### Supplementary Figure 5

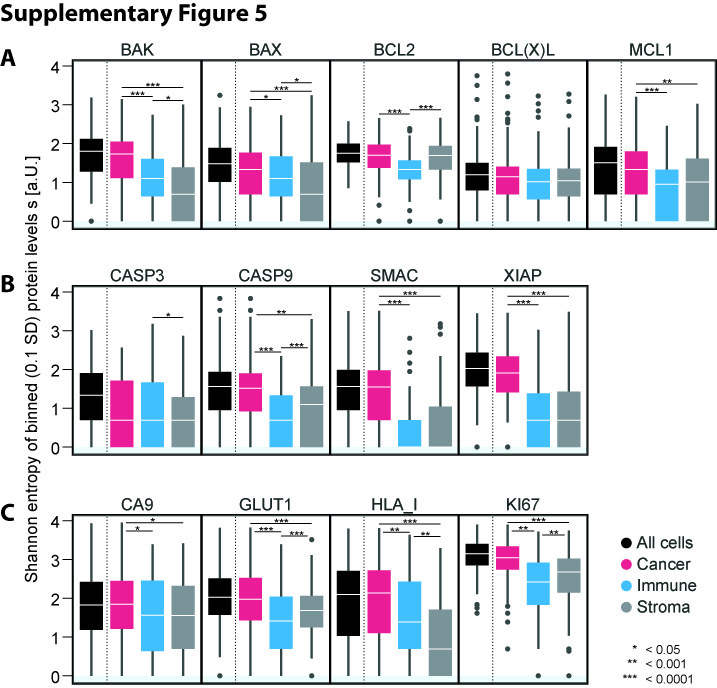

### Supplementary Figure 6

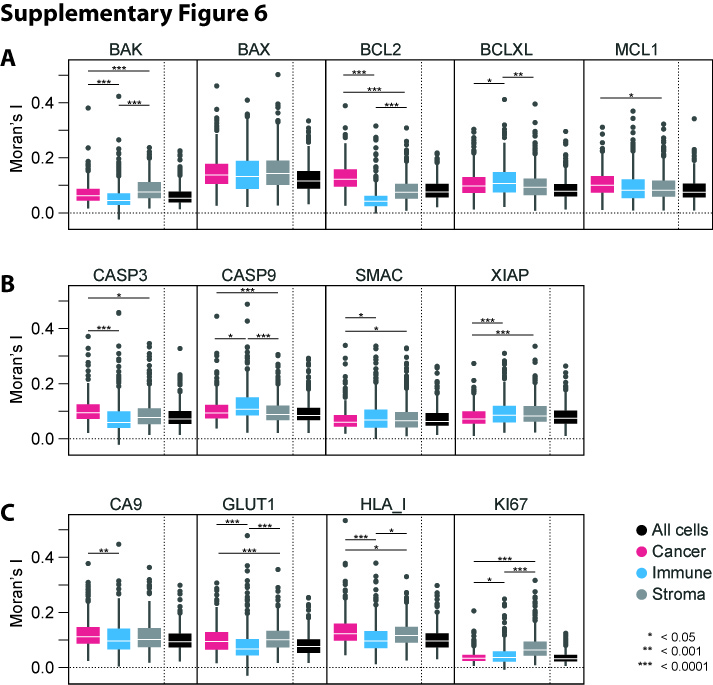
